# Paralytic Shellfish Toxin production in *Alexandrium minutum* (Dinophyceae): insights from omics integration using toxigenic and non-toxigenic recombinant progeny

**DOI:** 10.64898/2026.03.24.713948

**Authors:** Lou Mary, Julien Quere, Marie Latimier, Sébastien Artigaud, Hélène Hégaret, Mickael Le Gac, Damien Réveillon

## Abstract

Paralytic Shellfish Toxins (PSTs) are produced by certain species of cyanobacteria and dinoflagellates. Part of the PST biosynthetic pathway has been elucidated in cyanobacteria, and the implication of some *sxt* genes has been confirmed by experimental studies. Contrary to cyanobacteria, knowledge about PST biosynthesis in dinoflagellates is more limited and generally restricted to comparative studies with the cyanobacterial pathway. To investigate the specificity of the PST pathway in dinoflagellates, 16 toxic and non-toxic *A. minutum* strains from a recombinant cross were compared, without prior assumption on genes or metabolites involved in PST synthesis, using an integrative approach combining untargeted metabolomic and transcriptomic data. Among the 60 most distinguishing transcripts between toxic and non-toxic strains, only 3 *sxt* genes were present, *sxtA4, sxtG*, and *sxtI*. In contrast, non-*sxt* homologs were detected as highly discriminant between these two phenotypes. More specifically, *a phyH* homolog may act as the analog of *sxtS* found in cyanobacteria. Moreover, we identified four putative synthetic PST intermediates. Among these, Int-C’2, correlated with the toxic phenotype, whereas 3 others were detected in both toxic and non-toxic strains, suggesting that these strains may share some parts of the biosynthetic pathway. Finally, our results showed that PST biosynthesis in dinoflagellate results from the activity of *sxt* genes, acquired by horizontal gene transfer from cyanobacteria, as well as from other genes not acquired from cyanobacteria, such as *phyH*.

## 1. Introduction

Dinoflagellates are aquatic unicellular eukaryotes that proliferate primarily in marine ecosystems, from shallow estuaries to the open ocean, and can be responsible for harmful algal blooms around the globe (Anderson et al., 2012). These enigmatic organisms possess a multitude of genomic features that make them particularly difficult to study, such as huge genomes (Lajeunesse et al., 2005), permanently semi-condensed chromatin (Livolant and Bouligand, 1978), and reduced transcriptional regulation (Van Dolah et al., 2007). Dinoflagellates produce a wide variety of specialized metabolites, some of them with toxic properties, in a broad sense, having detrimental effects on their competitors (“allelochemicals” or “BECs (Bioactive extracellular compounds)”, Tillmann et al., 2009, Long et al. 2021), on marine macrofauna (ichthyotoxins, lysins, Hallegraeff et al., 2023, Place et al., 2024), and on humans (PSTs, Ritchie & Rogart, 1977 and other bioaccumulated toxins (Turner et al., 2021)).

Paralytic Shellfish Toxins (PSTs), which parent molecule is the saxitoxin (STX), are produced by certain species of cyanobacteria (Mahmood & Carmichael, 1986, Carmichael et al., 1997, Lagos et al., 1999) and dinoflagellates of the genera *Alexandrium* (Shimizu et al., 1977), *Pyrodinium* (Harada et al., 1982), *Centrodinium* (Shin et al., 2020) and *Gymnodinium* (Oshima et al., 1987). These soluble alkaloids can block voltage-dependent sodium channels in vertebrate cells (Ritchie and Rogart, 1977) and induce Paralytic Shellfish Poisoning (PSP) in humans (Kao, 1993) due to their bioaccumulation through the trophic food web, including in organisms consumed by humans, such as shellfish (Bricelj & Shumway, 1998). Blooms of PST-producing species may therefore have sanitary consequences and lead to commercial closures, with socio-economic impacts on shellfish farms and fisheries (Etheridge, 2010).

The underlying genetic basis of the PST biosynthesis was discovered in cyanobacteria, with the identification of a gene cluster, the *sxt* genes, responsible for the biosynthesis of saxitoxin and its analogs (Kellmann et al., 2008, Mihali et al., 2009, 2011). These genes consist of 14 essential “core genes” (Murray et al., 2011), *sxtA-I, sxtP-R, sxtS*, and *sxtU*, as well as tailoring genes *sxtL, N, X*, and regulators *sxtY, sxtZ* and *ompR* (Kellmann et al., 2008). So far, the biochemical function of only a few *sxt* genes in the biosynthetic pathway has been experimentally validated in cyanobacteria (e.g. *sxtA, sxtG, sxtH, sxtT*, and *sxtN*, Lukowski et al., 2018, 2019, 2020, Cullen et al., 2018, Chun et al., 2018).

Due to their genetic and physiological peculiarities, dinoflagellates represent a challenge for applying classical gene validation methods, explaining why the PST biosynthetic pathway is only partially elucidated in this taxon. In dinoflagellates, knowledge about the PST biosynthesis is thus primarily based on our understanding of the cyanobacterial pathway, as well as on comparative studies involving a small number of clones. For instance, blast search and function-based search of the cyanobacterial *sxt* genes against dinoflagellate transcriptomes are common procedures, leading to the discovery of hundreds of “*sxt* homologues” in these datasets (Hackett et al., 2013, Zhang et al., 2014, Muhammad et al. 2025). However, these putative *sxt* genes present different levels of homology with their cyanobacterial counterparts. Dinoflagellate *sxtA, sxtG* and *sxtB* are phylogenetically closely related to their cyanobacteria homologs and in-depth studies have been conducted on their role in PST synthesis in dinoflagellates (Stüken et al. 2011, Orr et al. 2013a, Murray et al. 2015, Kim et al. 2023, Cho et al., 2021, Cho et al., 2024). For other *sxt* genes such as *sxtD, sxtS* and *sxtU*, homology is solely based on domain similarity, leading to numerous putative homologs without close phylogenetic relationship to proteins from toxic cyanobacteria (Hackett et al., 2013) and their role in the PST biosynthesis pathway is still hypothetical (Akbar et al., 2020). In a previous study, we showed that, in *A. minutum*, a genomic region was associated with the ability to produce PSTs, and that some putative cyanobacterial *sxt* homologs are scattered throughout the genome and do not form a cluster (Mary et al., 2022), as observed in cyanobacteria (Kellmann et al., 2008). A similar observation was recently reported for *sxtA, sxtG* and *sxtB* in *A. catenella* (Kim et al., 2024). Dinoflagellates might exhibit specific features regarding PST synthesis (e.g. specific set of genes), which may have been overlooked by research based on similarity to *sxt* genes.

Finally, while studies focusing on the link between *sxt* genes and biosynthetic intermediates remain scarce, intermediates A’, C’2, 11-hydroxy-Int-C’2, E’, and the shunt product Cyclic-C’ have been detected in the PST-producing species *A. catenella* (Cho et al., 2016, 2019; Tsuchiya et al., 2014, 2016, 2017). Using HRMS (High Resolution Mass spectrometry) and synthesized putative precursors, the same research team was recently able to detect dd-doSTX in toxigenic *G. catenatum* and *A. pacificum* (group IV), as well as 12β-d-doSTX in toxigenic A. catenella and A. pacificum (group IV) (Hakamada et al., 2024). As for *A. minutum*, Int-A’, E’ and cyclic-C’ were reported, using an untargeted metabolomic approach (Brown et al., 2022). Altogether, these studies showed that LC-HRMS based untargeted metabolomics can be useful to identify, without a priori, metabolites that may be involved in PST biosynthesis.

The aim of this study was to move beyond the prevailing cyanobacteria-centred framework of PST biosynthesis in dinoflagellates. Previous investigations largely relied on homology to cyanobacterial *sxt* genes or on targeted searches for candidate metabolites (Hackett et al., 2013; Zhang et al., 2014; Muhammad et al., 2025). We implemented an integrative and untargeted approach, combining metabolomics and transcriptomics, on toxic and non-toxic *Alexandrium minutum* strains derived from a recombinant cross. Because PST production segregates among genetically related progeny, gene expression profiles and metabolites that co-segregate with toxicity can be identified independently of prior pathway assumptions. This design strengthens the inference of functional associations between gene expression, metabolite profiles, and PST production, thereby enabling the identification of candidate components of a dinoflagellate-specific biosynthetic framework.

## 2. Material and methods

### 2.1. Generation of F1 strains

Each F1 strain used is a recombinant clone obtained from a cross between a toxic strain (T2) and a non-toxic strain (NT2) of *A. minutum*, as previously described (Seveno et al., 2020, Mary et al., 2022). Briefly, the cross was performed by mixing 1 mL of each parental culture (5000 cells/mL) to produce resting cysts, which then underwent an 8-month dormancy period at 4°C. After 5 days, in an algal incubator at 18°C, clonal recombinant F1 strains were isolated by a series of dilutions. SNPs discriminating the two parental strains were used to verify that the isolated strains were clonal and recombinant (Mary et al., 2022).

### 2.2. Experimental design

The 16 *A. minutum* strains were chosen according to their previously assessed toxin profiles (Mary et al. 2022): the 2 parents (one toxic, T2, one non-toxic, NT2), and their 14 recombinant F1 strains (the toxic strains T, F, K, M, C, L, Y, W, and the non-toxic strains P, X, U, Q, N, B). Five replicates (named A, B, C, D, and E) of each strain were grown, and their growth started one day apart (all replicates A on day 1, replicates B on day 2, replicates C on day 3, D on day 4, and E on day 5). These 80 samples (16 strains x 5 replicates) were grown in 420 mL of K+Na_2_SiO_3_ medium (Keller et al., 1987), in an algal incubator, in a 12/12 L:D cycle at 18°C for 7 days. At the end of the 7 days, for each sample, 3×100 mL were pelleted to extract intracellular metabolites, RNA, and PSTs (see Mary et al. 2025 for methods). Sampling took place after 12 h of darkness followed by 1 h of light, so that the strains could readjust to the light before sampling.

### 2.3. Metabolite extraction and analysis

While PST profiles were obtained using dedicated extraction (including a solid phase extraction step) and HILIC-MS/MS procedures (see 2.2), our analytical strategy for metabolomics was aligned with our untargeted approach, and based on a single-step methanolic extraction (i.e. minimal pre-treatment), and a conventional LC-HRMS method.

Cell pellets were stored at −20 °C until extraction. They were extracted with methanol (Chromasolv, Honeywell, LC-MS grade) at a ratio of 700 µL for 1 million cells, in an ultrasonic bath for 15 min at 25 kHz (sweep mode). An empty tube (Eppendorf) was processed as cell pellets and accounted for a procedural blank (i.e. used to filter out contaminants). The qualitative control (QC) sample was prepared by pooling the same volume of all methanolic *A. minutum* extracts. Finally, aliquots were put into glass inserts (250 µL, Interchrom, Interchim) and stored at −80 °C until analysis.

Metabolomic profiles were acquired by liquid chromatography coupled to high resolution mass spectrometry (LC-HRMS) as in Gémin et al. (2020). The system was a UHPLC (1290 Infinity II, Agilent technologies, Santa Clara, CA, USA) coupled to a quadrupole-time of flight mass spectrometer (QTOF 6550, Agilent technologies, Santa Clara, CA, USA) equipped with a Dual Jet Stream ESI interface. Both positive and negative full scan modes were used over a mass-to-charge ratio (*m/z*) ranging from 100 to 1700. The only difference with Gémin et al. (2020) was that the injection volume used here was 10 µL. The batch was prepared as recommended by Dunn et al. (2011), by using a mix of 10 phycotoxins as the Standard Reference Material. The injection order was randomized and QCs were injected every 8 samples.

LC-HRMS raw data (file format .d) were converted to .mzXML format using MS-Convert (ProteoWizard 3.0) (Chambers et al., 2012) and pre-processed with the Workflow4Metabolomics 4.0 e-infrastructure (Guitton et al., 2017). Peak picking (with a filter on retention time 0-780 s), grouping, retention time correction and peak filling were performed with the “CentWave”, “PeakDensity”, “PeakGroups”, and “FillchromPeaks” algorithms. Annotation (isotopes, adducts) was conducted with the “CAMERA” algorithm (Kuhl et al., 2012). Intra-batch signal intensity drift was corrected by fitting a locally quadratic (loess) regression model to the QC values (Dunn et al., 2011; Van Der Kloet et al., 2009).

Three successive filtering steps using in-house scripts on R were applied to remove low intensity, noisy and redundant variables, as in Georges des Aulnois et al. (2020). Pre-processing of +MS and - MS data matrices led to 10279 and 3773 variables respectively, and 1290 and 713 variables remained after the filtrations, respectively. The two matrices were concatenated and log transformed before statistical analyses.

In an attempt to annotate the significant features revealed by the metabolomic approach, MS/MS spectra were obtained, by auto MS/MS as in Georges des Aulnois et al. (2019) and by targeted MS/MS (supplementary tables 1 and 2). All spectra were processed using GNPS (Wang et al., 2016), SIRIUS (Dührkop et al., 2019), and MoNa (MassBank of North America, http://mona.fiehnlab.ucdavis.edu). In the absence of standards, the putative annotation of biosynthetic intermediates was carried out by comparing the exact mass and MS/MS spectra obtained with those existing in the literature (Kellmann et al., 2008, Cho et al., 2019, Brown et al., 2022). The level of annotation obtained ranged from 5 (exact mass) to 2 (probable structure by comparison with literature/library spectrum), according to Schymanski et al., (2014). All annotation details (e.g. formula, mass error, putative identity, class prediction or MS/MS spectra) were provided in the supplementary tables 1-6. Acquisition and data processing were performed using MassHunter Workstation softwares (B.09 and B.07, Agilent). All data (HRMS, autoMS/MS and targetedMS/MS) were deposited on DATAREF (https://sextant.ifremer.fr/record/174d9201-7633-436c-ba56-c9f226134026/).

### 2.4. RNA extraction, sequencing and annotation

Cell pellets were lysed by sonication on ice (Seveno et al., 2020; Mary et al., 2022). RNA from each culture was extracted using a NucleoSpin TriPrep kit, from Macherey-Nagel, following the manufacturer protocol. RNA was stored at −80°C. Out of the 80 original samples, one RNA extraction failed.

Library preparation was performed using the Illumina TruSeq mRNA stranded kit starting from 0.5 µg of total RNA. Library quality and quantity were assessed on a Bioanalyzer (Agilent Technologies, Palo Alto, CA) using high-sensitivity DNA analysis chips. Paired-end sequencing (2×150bp) of the 79 libraries was performed on two lanes of Flowcell SP on Illumina NovaSeq6000 at the GeT-PlaGe France Genomics sequencing platform (Toulouse, France). Raw sequencing data were submitted to the ENA (PRJEB98293).

Reads were trimmed using Trimmomatic version 0.36 (Bolger et al., 2014) with the parameters LEADING 3, TRAILING 3, MINLEN 36. The adapter file used was TruSeq3-PE. A maximum of 2 mismatches for the seed, a minimum score of 30 for the extension, and a minimum adapter length of 2 were used. Finally, only paired forward and reverse reads were kept. The reads were then aligned using the bwa mem command in the bwa aligner version 0.7.15 (Li and Durbin, 2009) to the *A. minutum* reference transcriptome (Le Gac 2016), with default parameters. A coverage file was generated for each strain using Samtools version 1.4.1 (Danecek et al., 2021) and variant calling was performed on each file using Freebayes version 1.3.5 (Garrison and Marth, 2012), restricting the analysis to previously identified SNPs (Seveno et al., 2020).

For each sample, the identity of the strain was confirmed using a custom python script, with a threshold <1% difference on more than 400 000 previously identified SNP genotypes (Seveno et al. 2020, Mary et al. 2022). One of the 79 samples was discarded due to missing data. The raw count matrix initially containing the count values of 153222 genes was filtered to remove low counts genes (average read counts per sample <10), resulting in a matrix of 104432 transcripts for 78 samples.

RNA annotation has been described in Mary et al (2022). Briefly, the reference transcriptome was annotated based on the Uniprot/Swissprot database (The Uniprot consortium, 2020). A tblastn (Altschul et al., 1990) using cyanobacterial *sxt* genes was performed against the reference transcriptome. These annotations can be found in supplementary tables in Mary et al (2022). ORFs were generated from the transcriptome using Transdecoder and were searched against the HMM library pfam-A.hmm (Mistry et al., 2021) using HMMER program (Eddy, 2011).

### 2.5. Statistics

Two complementary approaches were used to determine which transcripts and metabolites are linked to PST production in toxic strains of *A. minutum*.

The first one is a classical method used for differential gene expression that was initially developed for microarray RNA analysis, and is implemented in the Limma R package (Ritchie et al., 2015). It will be hereafter referred to as « the linear model » and was used on the two separate datasets (transcripts and metabolites), using replicates as random effect. This first approach is not a traditional one for analyzing metabolomics datasets, for which multivariate models are established methods (Xi et al., 2014). Therefore, a second approach based on a multivariate model was used, and consisted in an integrated supervised sparse Partial Least Square Discriminant Analysis (sPLS-DA), performed on both datasets. This second method combines classical PLS-DA (dimension reduction and supervised classification) and the selection of a subset of independent variables with the greatest discriminating power between categories (“toxic” and “non-toxic” in this study) (Lê Cao et al., 2011). It was developed for biomarker discovery in omics studies (DIABLO, Singh et al., 2019), and is implemented in the Mixomics package (Rohart et al., 2017).

#### 2.5.1. Separate analysis using the two datasets (linear model)

For the RNA dataset, scale (library size) normalization was performed on the 104432 transcripts x 78 samples matrix, using the EdgeR R package (v.4.6.2, Chen et al., 2025). Count data were then transformed to log2 count per million using the voom function of the Limma R package (v.3.64.1), and linear model was applied accounting for strain-effect. For the metabolite dataset, the already log-transformed matrix of 2003 features x 78 samples was used for this analysis, and as for RNA, a linear model was applied, using replicates as a random effect.

#### 2.5.2. Data integration using sPLS-DA

##### 2.5.2.1. RNA matrix pre-processing

The 104432 transcript matrix was log normalized using the vst function of the Deseq2 package version 1.26.0 (Love et al., 2014) on R version 3.6.3, with a design taking into account the replicates. Homogeneity of expression between replicates was checked using a PCA and expression heatmap. As recommended in the Mixomics package’s vignette, transcripts displaying high gene expression variability between replicates were discarded by filtering out transcripts for which the Median Absolute Deviation (MAD) between replicates of a strain was > 0.25. To focus on transcripts displaying strong gene expression differences between strains, the MAD was then calculated for each transcript between all samples, and only transcripts with MAD >= 0.25 were retained. Finally, for each transcript, the median expression across replicates was calculated and retained. The final expression matrix corresponded to 16 strains x 4201 transcripts.

##### 2.5.2.2. Metabolite matrix pre-processing

As with the RNA matrix, the normalized metabolite matrix (initially 2003 features) was filtered by calculating the MAD, with a maximum threshold of 0.8 for MAD between replicates, and a threshold greater than 0.1 between samples, yielding a 16 strains x 1442 metabolites matrix. Thresholds are different from those used for RNA due to greater heterogeneity between replicates for this dataset.

##### 2.5.2.3. sPLA-DA parameters and cross validation

An integrated supervised sparse Partial Least Square Discriminant Analysis (sPLS-DA) “block” (i.e. coupling metabolomics and transcriptomics data), was performed using the two log-normalized RNA and metabolite matrices, and a Y matrix indicating the toxic or non-toxic group membership of each strain, using the package MixOmics (v.6.32.0). The parameters of the design matrix are 0.5 correlation between the RNA and metabolite datasets, and 1 correlation between the toxic/non-toxic matrix and the two RNA and metabolite datasets. Due to the small number of strains, the “Leave-one-out” cross validation (a special case of k-fold cross validation) procedure was used. The sPLS-DA model was tuned to find the adequate number of components and features of each dataset to minimize classification and balanced error rate (supplementary fig. 1A, 1B).

### 2.6. Blast and PhyH phylogenetic analysis

#### Blast

All candidate transcripts from the sPLS-DA analysis, overexpressed in toxic strains, were blasted against Eukprot Alveolate database V3 (Richter et al., 2022) with an e-value threshold of 1e-5. Emphasis was then put on transcripts matching sequences from other PST-producing dinoflagellates (*P. bahamense* and *G. catenatum*). In particular, the candidate transcript comp103275_c0_seq1 (see results), annotated as a Phytanoyl-CoA dioxygenase (*phyH*) shared the same function as the *sxtS* gene from PST-producing cyanobacteria, and was further investigated.

#### PhyH phylogenetic analysis

All *phyH* and *sxtS* sequences used were collected from GenBank and Eukprot V.3 databases, and are available in supplementary table 7. Dinoflagellate sequences from Eukprot database matching *PhyH* transcript (see results) were aligned with cyanobacterial SxtS and other Phytanoyl-CoA dioxygenase sequences in order to assess if our *PhyH* transcript was inherited from cyanobacteria. Alignment was realized using the Muscle algorithm in MEGA X version 10.1 (Kumar et al., 2018), with default parameters (Gap Open −2.9, Gap Extend 0.0, Hydrophobicity multiplier 1.20, Cluster method UPGMA, Min diag len 24), manually checked, and poorly aligned positions were excluded. The optimal substitution model was VT+G4 according to the BIC criterion, and was assessed using ModelFinder (Kalyaanamoorthy et al., 2017). Tree inference was computed using Maximum Likelihood and consensus tree was selected based on 1000 bootstrap trees using IQ-TREE (Nguyen et al., 2015). Ultrafast bootstrap (UFBoot, Hoang et al., 2018) and SH-aLRT (Guindon et al., 2010) were performed as branch support analysis. IQ-TREE web server (Trifinopoulos et al., 2016) was used to select the model and compute trees and branch support. PhyH sequences from Archaea were used as an outgroup. The active site (metal coordinating residues) of the PhyH sequences was inferred according to Clifton et al. (2006) and Schofield & McDonough (2007). Aligned amino acids in the vicinity of this active site were plotted for all dinoflagellates and PST-producing cyanobacteria (SxtS) using the R library ggseqlogo (Wagih, 2017).

## 3. Results

### 3.1. PST profiles

PST analogs were detected in 9 of the 16 strains using LC-MS/MS (Mary et al., 2025, the two studies were conducted at the same time on the same samples). Briefly, PSTs were detected in the parental strain T2 and 8 F1 strains: W, C, F, Y, K, L, M. The parent NT2 and F1 strains X, U, N, P, Q and B did not produce PSTs at detectable levels, and are considered as ‘non-toxic’ (fig. 1 A, B).

**Figure 1:**
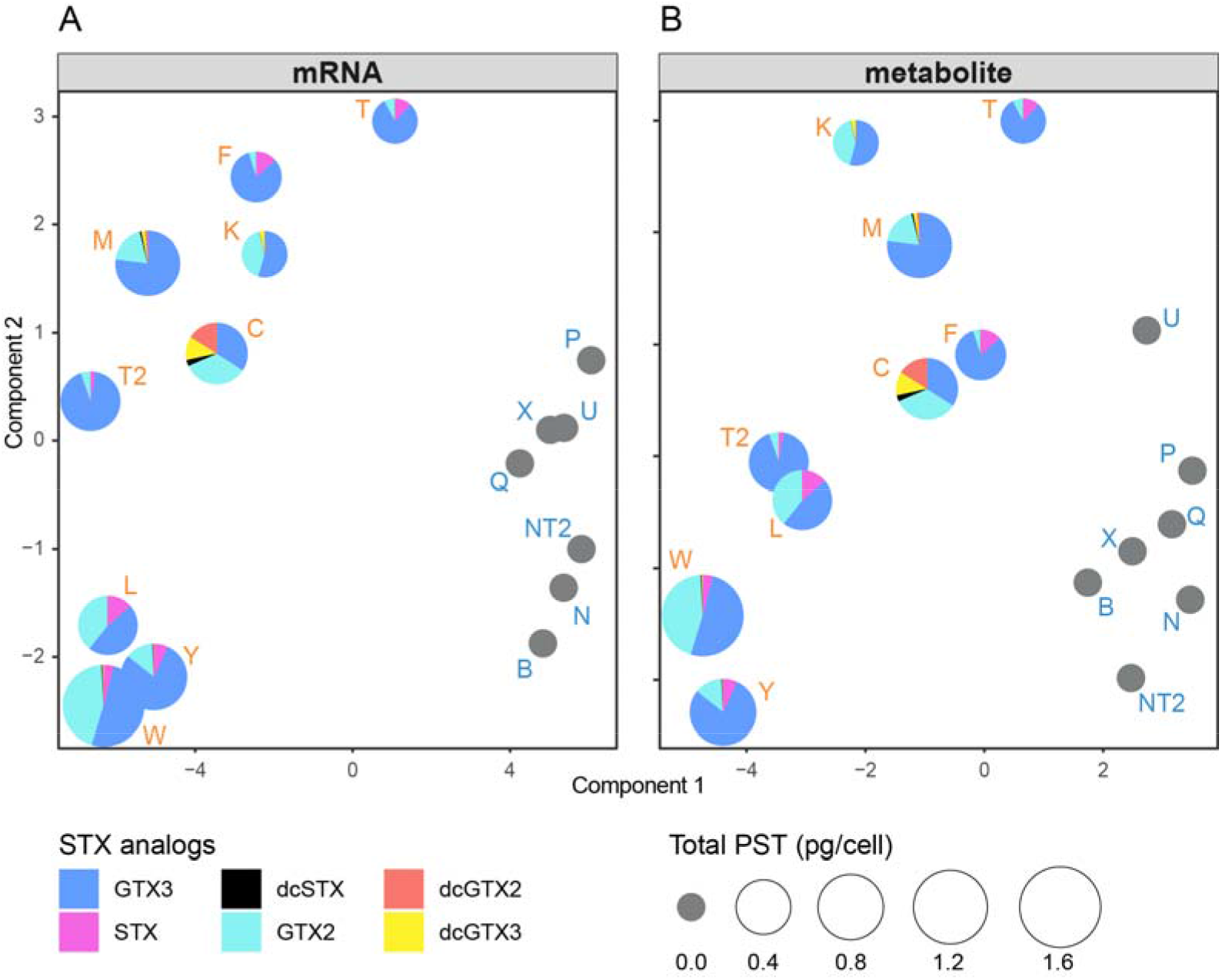
**A, B:** separation of strains after sPLS-DA analysis according to their toxicity based on 60 transcripts **(A)** and 27 metabolites **(B)**. PST profiles are indicated by pie-charts, size of the charts indicates total PST content per cell (pg/cell).

### 3.2. Genes and metabolites correlated with toxin production in *A. minutum*

#### 3.2.1. Transcripts and metabolites from both linear model and sPLS-DA

As described in 2.5., two approaches were used to analyze the two datasets: a linear model for each dataset using replicates, and a joint multivariate model using median values for each strain (sPLS-DA).

Using the sPLS-DA approach, from the initial dataset of 4201 transcripts and 1442 metabolites, 60 transcripts and 30 metabolites were retained as the most discriminating features between the 16 toxic and non-toxic strains (balanced error rate on component 1=0.055). Of the 60 differentially expressed transcripts, half were overexpressed in toxic strains (fig. 2A). Of the 30 metabolites selected by sPLS-DA, 3 were subsequently excluded as they corresponded to isotopes of non-significant features. Of the 27 remaining metabolites identified with this method, 12 were overexpressed in toxic strains (fig. 2B). While a PCA on the initial whole dataset did not show any separation between toxic and non-toxic strains (not shown), the sPLS-DA model allowed this discrimination on the first component (fig. 1 A, B). The discriminating transcripts and metabolites were ranked based on their contribution (i.e. importance) to the toxic/non-toxic phenotype (fig. 2A and 2B respectively).

**Figure 2:**
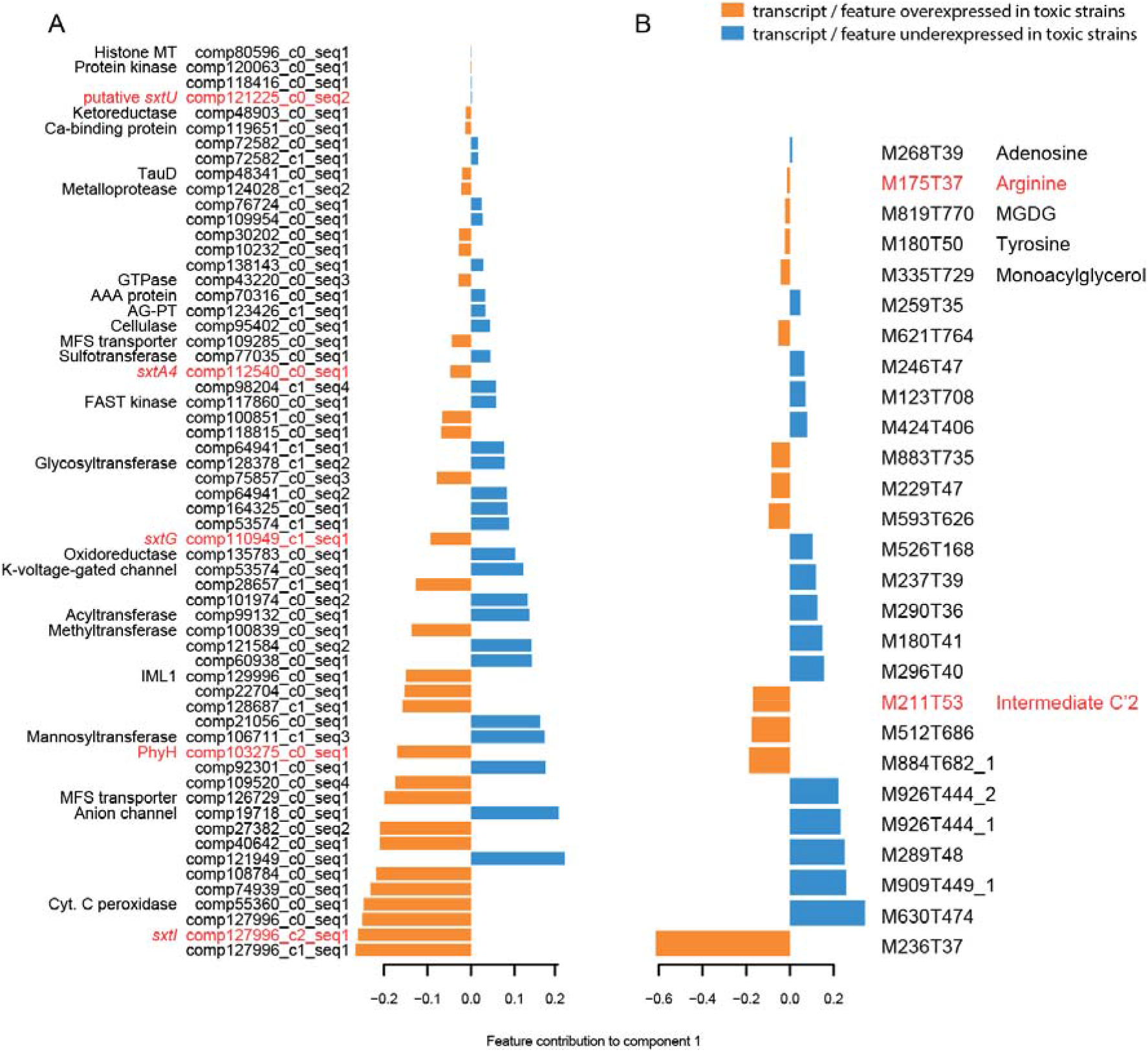
**A, B:** Contribution (i.e. importance) of each feature on the first component (blue bar: more expressed in non-toxic strains, orange bar: more expressed in toxic strains), for RNA **(A)** and metabolites **(B)**. Variable contributions on component 1 are ranked according to their importance from bottom to top. Transcripts and features in red are the ones known to be involved in cyanobacterial PST pathway (analogous or homologous). AG-PT: aminoglycoside phosphotransferase, Cyt-c: cytochrome-c, MFS: major facilitator superfamily (transporter), MT: methyltransferase, MGDG: monogalactosyldiacylglycerol, PhyH: phytanoyl-CoA dioxygenase, TauD: taurine catabolism dioxygenase.

Using the linear model approach, all but 6 of the 60 transcripts found with the sPLS-DA model showed adjusted p-value < 0.05 (supplementary fig. 2, 3, supplementary table 8), and all but 6 of the 27 metabolites showed adjusted p-value < 0.05 (supplementary fig. 4, 5, supplementary table 9).

Of the 60 transcripts identified with sPLS-DA, 27 were annotated (supplementary table 8). Of the 27 metabolites identified with sPLS-DA 6 were annotated with a level 2 of annotation, and a compound class prediction could be obtained for 16 features (level 3 of annotation, Schymanski et al., 2014, supplementary table 3).

#### 3.2.2. Metabolites contributing to the toxic phenotype

The metabolites contributing most to the toxic phenotype according to the sPLS-DA was by far M236T37 (|weight|=0.6) (fig. 2B), followed by M884T682_1, M512T686, and M211T53. This last feature was annotated as the PST biosynthetic intermediate Int-C’2 (level 2 of annotation, supplementary table 5). In the linear model, M236T37 was >4 times more expressed in toxic strains, M211T53 (Int-C’2), ~3 times more expressed, and M512T686, 2 times more expressed in toxic strains (adj. p-values = 2.5e-22, 1.1e-5 and 1.8e-4, respectively). Arginine (PST precursor) was selected by sPLS-DA and overexpressed in toxic strains. However, this feature did not meet the threshold in the linear model.

Three metabolite features linked to PST biosynthesis were initially present in the complete dataset but not retained because they were either filtered out during data processing or did not show sufficient variability between the two groups of toxic and non-toxic strains. The first one, M225T32, was annotated as the PST intermediate 12-β-d-doSTX (level 2 of identification, supplementary table 6). The second feature, M209T36, could match the shunt product of the PST pathway, Cyclic-C’, or another precursor, dd-doSTX (same exact mass, level 4 of identification, supplementary table 6). Finally, M241T32 might correspond to the STX precursors doSTX or 12-β-d-dcSTX (level 4 of identification, supplementary table 6). Normalized values for these features were compared between the toxic and the non-toxic group. The differences between the two groups were non-significant for the 3 features (p-values<0.05, supplementary fig. 6), using non parametric tests.

#### 3.2.3. Genes contributing to the toxic phenotype

Among the 68 transcripts putatively matching 12 cyanobacterial *sxt* genes present in the *A. minutum* transcriptome (Mary et al. 2022), only 3 were overexpressed in toxic strains in this study, and selected by sPLS-DA: *sxtI* (|weight|=0.26), *sxtG* (|weight|=0.09) and *sxtA4* (|weight|=0.05) (fig. 2A). A transcript annotated as phytanoyl-CoA dioxygenase (*PhyH*), was also identified as overexpressed in toxic strains. This transcript shares the same function as the *sxtS* gene in PST-producing cyanobacteria. Among the transcripts overexpressed in non-toxic strains, a *sxtU* putative homolog, was identified as a weak contributor to the non-toxic phenotype (|weight|=0.002). These transcripts were also found with the linear model: *PhyH* and *sxtI* were respectively 38 times and 19 times more expressed in toxic strains, while *sxtA4* and *sxtG* were 2x more expressed. Putative *sxtU* was 5x more expressed in non-toxic strains.

25 overexpressed sequences in toxic strains matched at least one sequence in the Eukprot Alveolate database. For 9 of them, the first hits were sequences from PST producing dinoflagellates *P. bahamense, G. catenatum*, and other *Alexandrium* species. Among them were *sxtI, sxtG, sxtA4, PhyH*, a transcript with a TauD domain and a protein annotated as a protein kinase A (supplementary fig. 7). In addition to *sxtI*, the two most discriminating transcripts (comp127996_c0_seq1 and comp127996_c1_seq1) are derived from the same component during *A. minutum* transcriptome assembly (Le Gac et al., 2016). One of them, comp127996_c0_seq1, matched some sequences of PST-producing dinoflagellates also present in *sxtI* hits. Finally, a transcript belonging to the membrane transport major facilitator superfamily (MFS), matched *P. bahamense* and other *Alexandrium sequences* (but no *G. catenatum* sequences) (supplementary fig. 8). The remaining 16 sequences with a hit in the Eukprot Alveolate database matched PST-producing dinoflagellates as well as non-toxic ones (not shown).

##### 3.2.3.1. Investigating SxtS analog PhyH

No closely related homolog of *sxtS* has been identified in dinoflagellate transcriptomes. However, the cyanobacteria SxtS share a conserved domain with PhyH (Hackett et al. 2013). As a result, the *phyH* transcript over-expressed in the toxic strains could be an analog of the cyanobacterial *sxtS*. To explore this hypothesis a phylogenetic approach was used. The phylogenetic tree built for the PhyH amino-acid sequences (fig. 3) showed that dinoflagellate PhyH sequences clustered together in a well-supported clade (bootstrap (UFBoot) value=100). Inside this clade, two sister groups were present: one, named Dinoflagellate PhyH1, containing our PhyH sequence together with only PST-producing dinoflagellate PhyH sequences (bootstrap (UFBoot) value=100), and the other clade, named dinoflagellate PhyH2, with the rest of the dinoflagellate PhyH sequences (including sequences from PST-producing and non-PST-producing dinoflagellates). In this last clade, internal nodes were less well supported (bootstraps (UFBoot) < 95) and relationships between species for this protein did not reflect the ones established for dinoflagellates in general. The whole dinoflagellate clade was a sister group (with 96 bootstrap (UFBoot) support) to a cyanobacteria clade, which included SxtS sequences from PST-producing cyanobacteria and other cyanobacterial *phyH* sequences.

**Figure 3:**
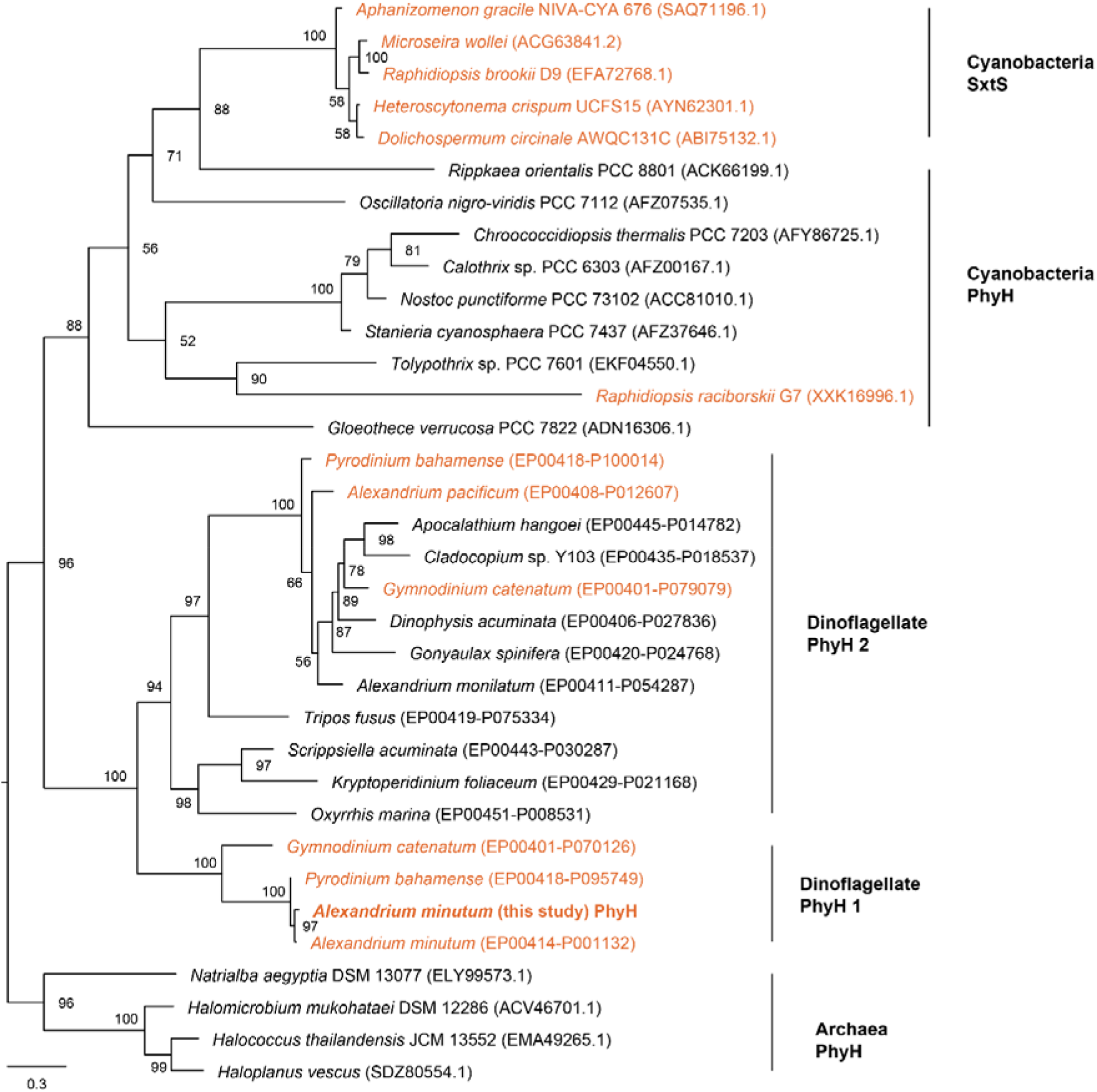
PhyH phylogenetic tree, inferred from 34 sequences of 180 amino-acid sites. The tree was generated using maximum likelihood, computed with IQTREE, using a model of substitution VT+G4. The consensus tree was built from 1000 bootstrap trees. Support values on nodes are IQTREE UFBoot2. Orange is for known PST-producing species.

An alignment of amino acids at the active site of dinoflagellate PhyH showed conserved ‘WHQ/ND’ residues for both PST-producing and non-PST-producing species (fig. 4 A, B, residue relative positions 2-5), and the same pattern was observed for the cyanobacterial SxtS used to construct the phylogenetic tree (fig. 4C). The sequence downstream of the active site is composed of more polar and acidic amino acids for PST-producing dinoflagellates than for non-PST-producing dinoflagellates (fig. 4 A, B, residues 6-24), whereas cyanobacterial SxtS downstream sequence is mostly constituted of hydrophobic residues.

**Figure 4:**
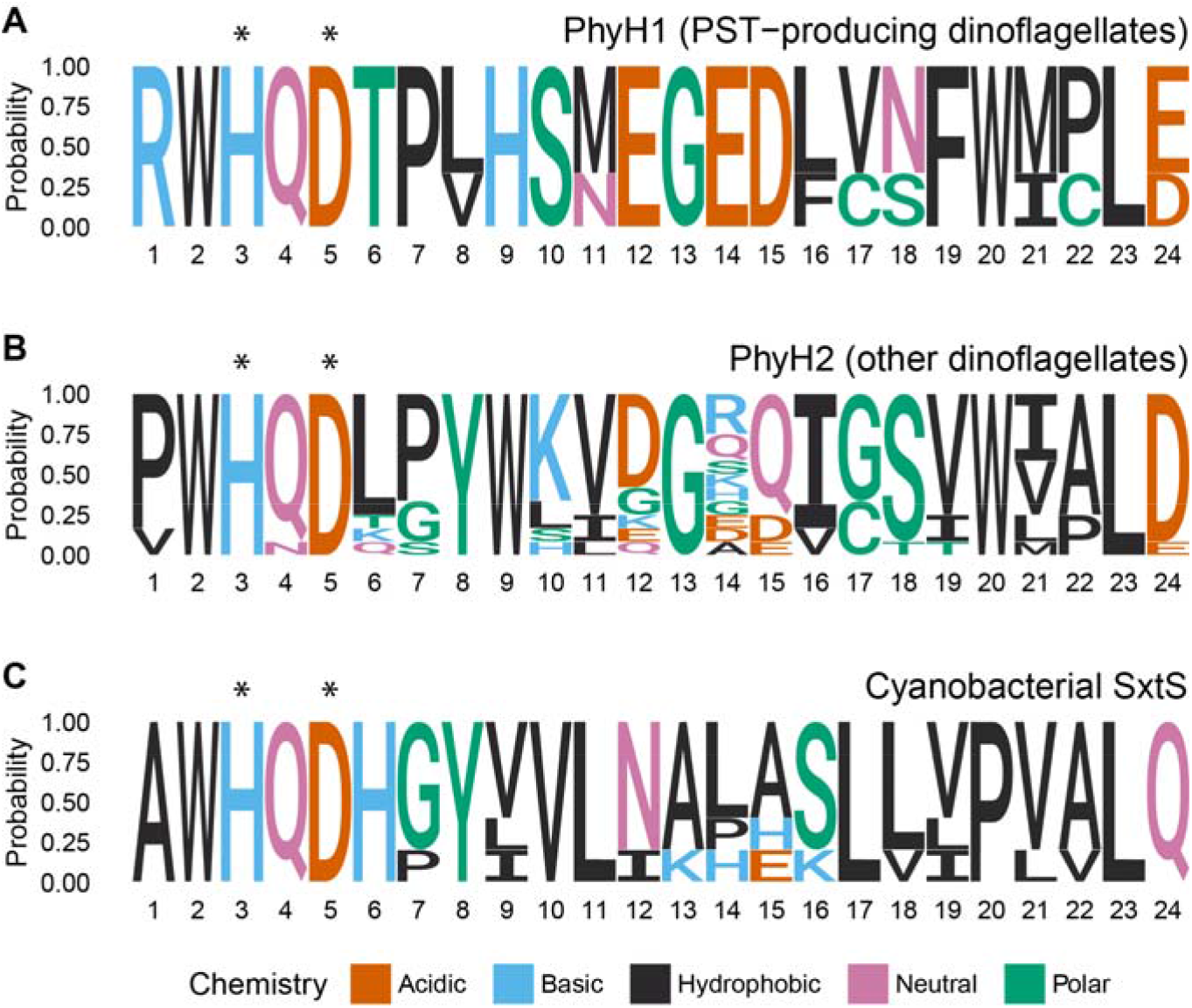
PhyH consensus protein sequence at the vicinity of the active site. The x-axis is the relative position of the amino acids, the y-axis is the probability to find them at a given position, based on the alignment of PhyH sequences. The two amino acids constitutive of the active site (inferred from Clifton et al. (2006) are indicated by stars (relative positions 3 and 5). **(A)** Consensus sequence for the PST-producing dinoflagellates (PhyH1, *A. minutum, P. bahamense*, and *G. catenatum*), with the “SN/MEGED” motif which seemed to be conserved, **(B)** Consensus sequence for the other dinoflagellates (PhyH2, PST-producing and non-PST-producing), **(C)** Consensus sequence for the PST-producing cyanobacterial PhyH SxtS sequences.

## 4. DISCUSSION

This study was designed to identify the gene expression profiles and metabolites that co-segregate with toxicity in *A. minutum* strains derived from a recombinant cross. This approach allowed the identification of candidate components of dinoflagellate-specific PST biosynthesis, independently of prior pathway assumptions. This was achieved by using an untargeted metabolomic approach and an untargeted transcriptomic approach. On the 12 putative *sxt* homologs previously found in the *A. minutum* transcriptome (Mary et al., 2022), only 4 were differentially expressed between toxic and non-toxic strains (*sxtI, sxtG, sxtA*, and *sxtU*), along with other non-*sxt* transcripts (comp127796_c0_seq1, comp100839_c0_seq1 (methyltransferase), and comp123426_c1_seq1 (phosphotransferase)). Furthermore, our method allowed us to detect a *phyH* transcript analogous but not homologous to *sxtS*, that might have an important role in the *A. minutum* PST pathway. Finally, five metabolites related to PST biosynthesis were identified, among which only two (Arginine and Int-C’2) were more expressed in toxic strains. These specificities are discussed and integrated in a comprehensive overview of the *A. minutum* PST pathway.

### First steps of the biosynthetic pathway (box 1, fig. 5)

**Figure 5:**
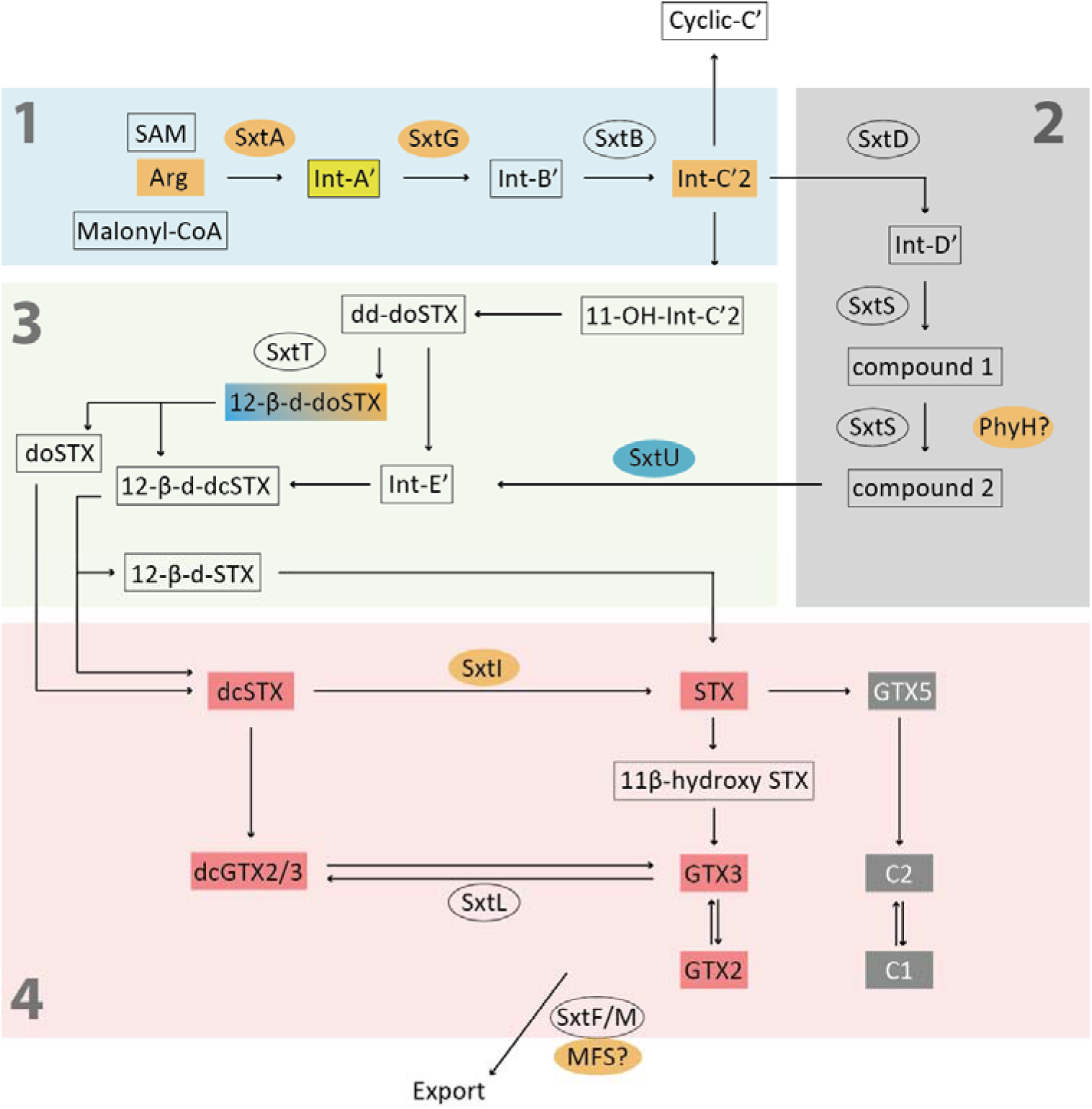
Putative PST biosynthetic pathway proposed for *A. minutum*, based on a review of the literature on PST-producing cyanobacteria and dinoflagellates (Kellmann et al., 2008, Mihali et al., 2011, Tsuchiya et al., 2014, 2017, Lukowski et al., 2018, 2019, 2020, Cho et al., 2019, 2023, Brown et al., 2022, Hakamada et al., 2024), as well as the toxin profiles observed in the cross studied here and previously (Mary et al., 2022). Box 1 represents steps of the pathway which are most certainly present in *A. minutum*. Box 2 represents a part of the initial pathway proposed by Kellmann et al. (2008), including two unnamed compounds, for which synthesis would be catalyzed by SxtS in cyanobacteria. Box 3 represents another route proposed by Hakamada et al. (2024), containing experimentally proven biosynthetic intermediates among which only 12-β-d-doSTX could be clearly identified in the present study with similar expression toxic and non-toxic strains (orange and blue). Box 4 represents the transformation of the different STX. Genes and metabolites overexpressed in toxic strains are shown in orange. Int-A’, in yellow, was identified in *A. minutum* in another study (Brown et al., 2022). Features overexpressed in non-toxic strains appear in blue. The *sxtS* analog PhyH and the transporter MFS are shown with question marks at points of the pathway where they might be involved (see discussion). The STX analogs in red are those identified in the present study. The biosynthetic route from STX to C-toxins was elucidated in cyanobacteria (Lukowski et al., 2019).

The moderate overexpression of the *sxtA4* gene - involved in the first step of the PST biosynthetic pathway in cyanobacteria (Kellmann et al., 2008) - in the toxic strains in this study was not identified in our previous one, where more strains were analyzed (Mary et al., 2022). This gene is considered as a good marker of toxicity in dinoflagellates (Murray et al., 2015, Suikkanen et al., 2013, Stüken et al., 2011, Geffroy et al., 2021), being responsible for Claisen condensation on the acetyl-ACP with arginine, to form the first Int-A’ intermediate (Kellmann et al., 2008, Tsuchiya et al., 2014, Chun and Narayan, 2019) (fig. 5). The latter is an ethyl ketone with an [M+H]^+^ *m/z* value of 187.1553 (Tsuchiya et al., 2014), produced from arginine (*m/z* 175.1190). In our study, while arginine was identified among the metabolites contributing to the toxic phenotype, no metabolite corresponded to Int-A’. Nevertheless, this intermediate has already been identified in a toxin-producing *A. minutum* strain (Brown et al., 2022). Given that we identified the precursor required for the Int-A’ synthesis, Arginine, and the intermediate following Int-A’ in the biosynthetic pathway (Int-C’), it is reasonable to assume that *A. minutum* is able to synthesize Int-A’. The reason why it is not detected here may be linked to a high Int-A’ turnover (see after) or to a lack of sensitivity of the LC-HRMS approach.

In the present study, the *sxtG* gene, overexpressed in toxic strains, was the second most differentiating *sxt* gene, after *sxtI*. This gene was found in the transcriptomes of several PST-producing dinoflagellate species belonging to the genera *Alexandrium* (Stüken et al., 2011, Orr et al., 2013a) *Pyrodinium* and *Gymnodinium* (Hackett et al., 2013, Murray et al., 2015) and does not appear to be present in dinoflagellates that do not produce PSTs (Murray et al., 2015), other than those of the genus *Alexandrium* (Orr et al., 2013a). The expression of *sxtG* seems to be correlated with toxin production in *A. catenella* (Kim et al., 2021) and *A. minutum* (Hii et al., 2016), although one study showed that the amount of *sxtG* mRNA in *A. minutum* did not correlate with the amount of intracellular toxins, leading the authors to suggest the involvement of post-transcriptional mechanisms in toxin production (Perini et al., 2014). Within the PST biosynthetic pathway, *sxtG* encodes an amidinotransferase that is thought to act on the first intermediate, Int-A’, resulting in the formation of an intermediate B’ (*m/z* 229.17), that spontaneously cyclized to form the intermediate C’2 (*m/z* 211.1666) (Tsuchiya et al., 2014, 2017, Lukowski et al., 2020). This metabolite was overexpressed in toxic strains in the present study, and coupled with the overexpression of *sxtG*, we assume the existence of this biosynthetic step in *A. minutum* (fig. 5).

Int-C’2 is partially converted to a shunt product of the biosynthetic pathway, Cyclic-C’ (*m/z* 209.1509), which is mostly excreted outside the cell (Tsuchiya et al., 2016). This by-product was detected in the PST-producing organisms *Dolichospermum circinale* (previously named *Anabaena circinalis*) and *A. catenella*, and was absent in their non-toxic counterparts (Tsuchiya et al., 2015, Cho et al., 2016). It was also detected in *A. minutum* (Brown et al. 2022). In our dataset, a feature could correspond to Cyclic-C’, M209T36_2 (annotation level 4), but was removed prior to analysis because it was showing no significant variability between toxic and non-toxic strains. Altogether these results suggest that Cyclic-C’ might be present in *A. minutum*.

### From Int-C’2 to STX analogs (boxes 2-4, fig. 5)

The next steps of the pathway are still open to discussion in dinoflagellates. Initially, in cyanobacteria, Kellmann et al. (2008) had hypothesized the formation of an intermediate D’ from C’, leading to the formation of int-E’ (box 2 in fig. 5). The *sxtS* gene, together with *sxtU* and *sxtD*, could be responsible for the formation of the ring structure between the C’ and E’ intermediates (Kellmann et al., 2008), and belongs to the core cyanobacterial set of *sxt* genes (Murray et al. 2011). Cyanobacterial SxtS bears a phytanoy-CoA dioxygenase domain (PhyH), and its function in the STX synthesis pathway would be linked to this domain. This domain is used in homology search to identify putative *sxtS* homologs in dinoflagellate transcriptomes, leading to a large number of putative *sxtS* candidates (Hackett et al 2012, Muhammad et al. 2025). The role of PhyH/SxtS in STX biosynthesis remains poorly understood in dinoflagellates, but the *phyH* homolog more expressed in the toxic strains of the present study is a good candidate to be analogous to the phytanoyl-CoA dioxygenase function of the cyanobacteria *sxtS* gene. This gene was one of the strongest contributors of the difference between toxic and non-toxic strains (38-times more expressed in toxic strains). Furthermore, two paralogs seemed to exist for this gene in dinoflagellates, among which one, *PhyH1*, seemed to be specific to PST-producing dinoflagellates, whereas non-PST-producing dinoflagellates only possessed the other paralog, *PhyH2*. These paralogs seemed to share the same active site as in cyanobacteria SxtS, and could therefore be functional. The PhyH gene phylogeny also indicated an absence of horizontal gene transfer from cyanobacteria to dinoflagellates. Furthermore, PST-producing dinoflagellate PhyH1 shared conserved sequence patterns in the area of the active site, different from SxtS and PhyH2. Altogether our results suggested that PhyH could have a similar function to SxtS in the PST biosynthetic pathway in *A. minutum* (box 2, fig. 5), but do not seem to have a cyanobacterial origin.

Alternatively to Kellmann’s hypothesis, recent studies suggested that part of Int-C’2 pool would be converted to 11-OH-Int C’2 (m/z 227.1615) (box 3, fig. 5), shown to be present in toxic *A. catenella* (Cho et al., 2019). This intermediate was neither detected in our dataset nor referenced in the *A. minutum* literature. This step of the pathway cannot therefore be confirmed for this species. The 11-OH-Int-C’2 would then be converted to dd-doSTX, itself transformed to Int-E’ or to 12-β-deoxy-decarbamoyloxySTX (12-β-d-doSTX) by *sxtT* (box 3, fig. 5) (Hakamada et al., 2024). This last metabolite was identified in our dataset without any difference between toxic and non-toxic strains (supplementary fig. 6). Furthermore, putative homologs of *sxtT* were present in *A. minutum* transcriptome (Mary et al., 2022), but they were not differentially expressed. Therefore, if this step is present in *A. minutum*, its ability to differentiate toxic and non-toxic strains is still an open question for this species. While 12-β-d-doSTX was confidently identified here (by comparison with the MS/MS spectrum in Hakamada et al. (2024) and the presence of the discriminating ion at *m/z* 122.07), the presence of Int-E’ cannot be totally excluded in *A. minutum*. Indeed, both intermediates can coexist in the same strain of *A. catenella* (Hakamada et al., 2024) and Int-E’ had previously been identified in *A. minutum* (Brown et al., 2022) but before the first report of 12-β-d-doSTX (Hakamada et al 2024).

Finally, two intermediates with the same mass (*m/z* 241.1407) could be synthesized from 12-β-d-doSTX: 12-beta-deoxy-dcSTX (shown to be present in D. circinale and A. catenella, Cho et al., 2015) and doSTX (present in D. circinale, Hakamada et al., 2024). As stated before, 12-beta-deoxy-dcSTX could have been present in our initial dataset, but was not identified as differentially expressed between toxic and non-toxic *A. minutum* strains. Whether the lack of differential expression could be due to a high turnover should be further assessed, as a low abundance does not necessarily reflect a reduced synthesis. Finally, it is worth noting that the cyanobacterial one-step precursor of STX, 12-beta-deoxySTX (12-β-d-STX) (Lukowski et al. 2018), detected in *A. pacificum* (Akamatsu et al. 2022), was neither detected in our dataset nor in previous *A. minutum* studies.

The *sxtU* gene is the least differentiating of the four *sxt* genes in this study. *sxtU* was genetically predicted by Kellmann et al. (2008), and is thought to act on the aldehyde moiety of an STX precursor, forming the Int-E’ intermediate. Surprisingly, its expression appeared to be slightly higher in the non-toxic strains. As STX intermediates synthesized between Int-C’2 and dcSTX were found to be present in non-toxic strains, one hypothesis could be that these strains may have retained parts of the PST pathway. This would not necessarily contradict previous studies focusing on the early steps, as perhaps these non-toxic strains would have lost their toxicity at later stages in the biosynthetic pathway. Wang et al. (2020) suggested that *sxt* genes are shared by some non-PST producing strains and might have other functions as well. The overexpression of *sxtU* could have pointed to that direction, however, a large number (37) of potential *sxtU* homologs were previously identified in *A. minutum* transcriptome (Mary et al., 2022), displaying relatively low homology with cyanobacterial *sxtU* compared to genes like *sxtA4* or *sxtG*. This may suggest that these genes are not true *sxtU* cyanobacterial homologs.

The annotation of the above-mentioned features and the *sxtU* gene would need to be validated, which will help to confirm the steps pictured in box 3, fig. 5 (Hakamada et al., 2024) for *A. minutum*, since, for now, only 12-β-d-doSTX appeared to be clearly present in this species.

Finally, the *sxtI* gene expression correlated very strongly with the toxic phenotype in this study. This gene encodes a carbamoyltransferase (Kellmann and Neilan, 2007) that enables the transfer of a carbamoyl group to form carbamoylated analogs such as GTXs and STX, which dominate the PST profiles of the strains used in the present study. Kellmann et al., (2008) were the first to describe this gene, as present in all the cyanobacterial PST-producing strains. It was then shown on a strain of the cyanobacteria *Microseira wollei* producing no carbamoyl analogs (e.g. STX, GTX toxins, and C1/2 toxins in fig. 5) that this particular toxin profile was probably due to a truncation of the *sxtI* gene (Mihali et al., 2011). The authors postulated that in this strain, Int-E’ would give dcSTX, under the action of *sxtV,W,H/T* (Mihali et al., 2011) (this hypothesis is not represented on fig. 5 to facilitate understanding). The last part of the pathway (box 4, fig. 5) was partly hypothesized (first part, Mihali et al. 2011), and partly elucidated in cyanobacteria (second part (from STX to GTX 2/3 and C1/2), Lukowski et al. 2019), and was coherent with the toxin profile of our *A. minutum* strains from different crosses (this study and Mary et al., 2022). We therefore hypothesize that SxtI catalyzes the carbamoylation of dcSTX, forming STX in *A. minutum*.

### Candidate genes and metabolites

Other candidate transcripts selected by the analysis may play a role in the formation of PST in *A. minutum*, but the very low level of transcriptome annotations available for dinoflagellates (Stephens et al., 2018) hampered the validation of these candidates. Among the annotated transcripts, the comp126729_c0_seq1 transcript encoded a protein belonging to the Major Facilitator Superfamily (MFS). This family of proteins are transporters present in all classes of organisms and can transport a wide variety of compounds, including metabolites and drugs (Pao et al., 1998). While studies have shown the presence of some members of this family of transporters in dinoflagellates (Dagenais Bellefeuille & Morse 2016, González-Pech et al., 2017, Abassi and Ki, 2022, Zhuang et al., 2015), none have been linked to toxin production. Among the *sxt* genes, transporters belonging to another protein family, the MATE (Multidrug and Toxic compound Extrusion) family, have been identified (*sxtF* and *sxtM*, Kellmann et al., 2008). Neither of these two *sxt* genes was identified in *A. minutum* transcriptome, but other means of PST export could coexist within toxic *A. minutum* strains, as shown in the toxic cyanobacteria *M. wollei*, as a possible adaptation of transport mechanisms (Mihali et al., 2011). Finally, a transcript encoding a TauD domain was a weak contributor to the toxic phenotype. This domain was already identified in a previous study investigating genes differentially expressed between toxic and non-toxic strains of *A. minutum* (Yang et al., 2010), and homologs might be present in other PST-producing dinoflagellates. In addition, this domain was found encoded at the N-terminal position of the *sxtA* gene in *A. pacificum* (Bui et al., 2022). Altogether, these results suggest that the TauD domain may play a role in the PST biosynthetic pathway in *Alexandrium* species.

The present study showed that the feature M236T37 strongly correlated with the toxic phenotype (i.e. no detection in non-toxic strains), raising questions about its role in toxin synthesis. However, the putative mass and formula did not seem to be consistent with an STX precursor. Furthermore, M236T37 may be structurally related to M296T40, overexpressed in non-toxic strains (i.e. they shared similar ions in their MS/MS spectra, see supplementary table 4), which is probably not associated with STX intermediates either (based on its putative imidazopyridine chemical class). A plausible explanation is that M236T37 participates in a metabolic pathway unrelated to PST biosynthesis. Its co-segregation with the ability to produce PST could be due to physical linkage between the locus encoding for the synthesis of M236T37 and the locus previously identified as controlling PST production (Mary et al., 2022). Under this scenario, the overexpression of M236T37 observed in toxic strains would likely be confined to the present family of strains and would not represent a general characteristic of *A. minutum*.

### Implications in the PST biosynthetic pathway

Our results, together with the literature covering PST synthesis in dinoflagellates, showed that the PST biosynthetic pathway in *A. minutum* seems to be a patchwork consisting of genes from different origins. The first part (box 1, fig. 5) seems to be composed of “true functional homologs” of cyanobacterial *sxt* genes (as stated by Orr et al. 2013b). Our comparative approach has highlighted a *phyH* transcript analogous to *sxtS*, along with other transcripts which functions could be implicated in the PST pathway. The next steps of the pathway might thus not all be catalyzed by homologs of cyanobacterial *sxt* genes evolutionary acquired by horizontal gene transfer, but by other genes already present in dinoflagellates, like *phyH*.

Our metabolomic approach, based on a single-step methanolic extraction, provides wide metabolite coverage (Cajka & Fiehn, 2016). While the methanol extraction may have been suboptimal for highly polar metabolites such as PSTs, possibly explaining their absence from the metabolomic dataset, we were able to detect two biosynthetic intermediates as well as two additional putative intermediates (see 3.2.2). These metabolites, found in PST-producing dinoflagellates and cyanobacteria (Hakamada et al. 2024) (box 3, fig. 5), were present in toxic as well as non-toxic *A. minutum* strains used in this study. The presence of intermediates in non-toxic strains was unexpected, but indicates that non-toxic strains might have retained parts of the PST biosynthetic pathway. Recent advances have shown that the metabolic dynamics operating in the PST biosynthetic pathway may be more complex than previously anticipated, with varying intermediate pool sizes, implicating de novo synthesis, but also synthesis from previously stocked amino acid or intermediates (Cho et al., 2023). To go further, and clarify the biosynthesis of PSTs in *A. minutum*, more quantitative analytical methods should be used (Cho et al., 2015, 2016, 2019, Hakamada et al., 2024), along with genetic analyses. Overall, this study has shown that the toxin biosynthetic pathway in *A. minutum* appears to involve previously predicted intermediates and genes, as well as a number of new candidates, genes and metabolites, and offers an opening for future research, such as experimental confirmation of their involvement in the toxic phenotype.

## Abbreviations

PSTs: Paralytic Shellfish Toxins
STX: saxitoxin
GTX: gonyautoxin
dcSTX: decarbamoyl-saxitoxin
dcGTX: decarbamoyl-gonyautoxin
Int-A’: intermediate A’
Int-C’2: intermediate C’2
Int-E’: intermediate E’
dd-doSTX: 12,12-dideoxy-decarbamoyloxySTX
12-β-d-doSTX: 12β-deoxy-doSTX
MAD: Median Absolute Deviation

## Acknowledgements

Sequencing has been done at the GeT-PlaGe GenoToul sequencing platform (Toulouse, France). We thank the Ifremer RIC informatic and sebimer bio-informatic teams for support.

## CRediT

Conceptualization: LM, SA, HH, MLG, DR

Methodology: All

Validation:

Formal analysis: LM, JQ, ML, MLG, DR

Investigation: LM, JQ, ML, MLG, DR

Resources:

Data Curation:

Writing - Original Draft: LM

Writing - Review & Editing: All

Visualization: LM

Supervision: SA, HH, MLG, DR

Project administration:

Funding acquisition: SA, HH, MLG, DR

