## Supplementary figures for "Paralytic Shellfish Toxin production in *Alexandrium minutum* (Dinophyceae): insights from omics integration using toxigenic and non-toxigenic recombinant progeny"

**Supplementary Figures and legends**

**Supplementary fig. 1:** **(A)** Classification error rate as a function of the number of components during sPLS-DA model tuning. Centroids distances and balanced error rate (BER) were the better variables to check considering our unbalanced design, according to Mixomics vignette **(B)** Number of selected features for each block (RNA-metabolites) minimizing balanced error rate (minimum BER=0.05) for sPLS-DA model.

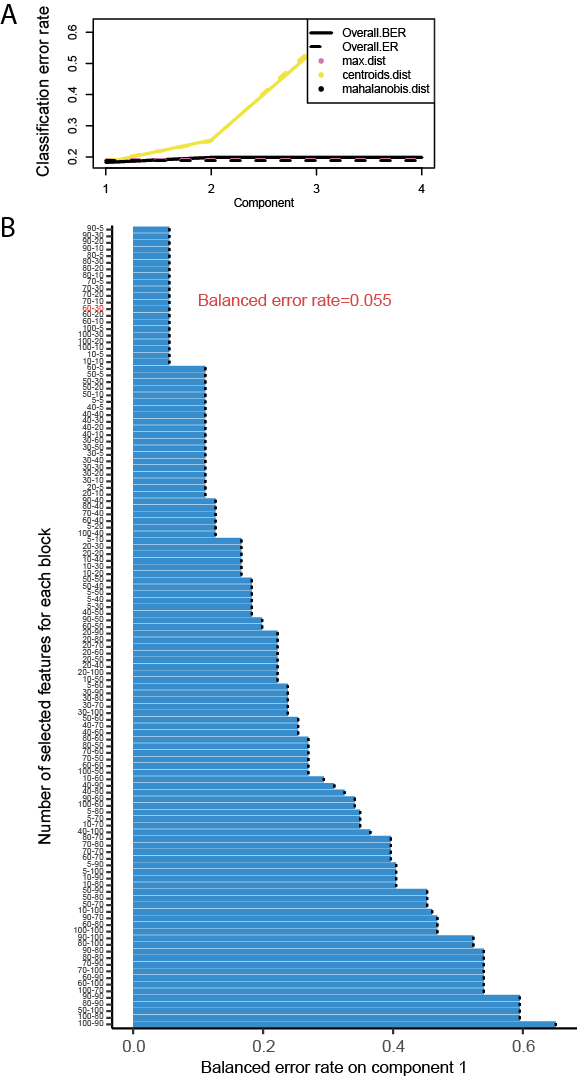

**Supplementary fig. 2:** Expression matrix computed with the linear model, showing overexpressed transcripts in **toxic** strains (all of the transcripts could not be represented, thus the results were filtered with a p-value threshold of 0.001). The blue to red scale indicates log2 fold change values between -2 (blue) and 2 (red) (i.e. Blue = underexpressed, red = overexpressed). Log fold change has been set between -2 and 2 for visualization purposes.

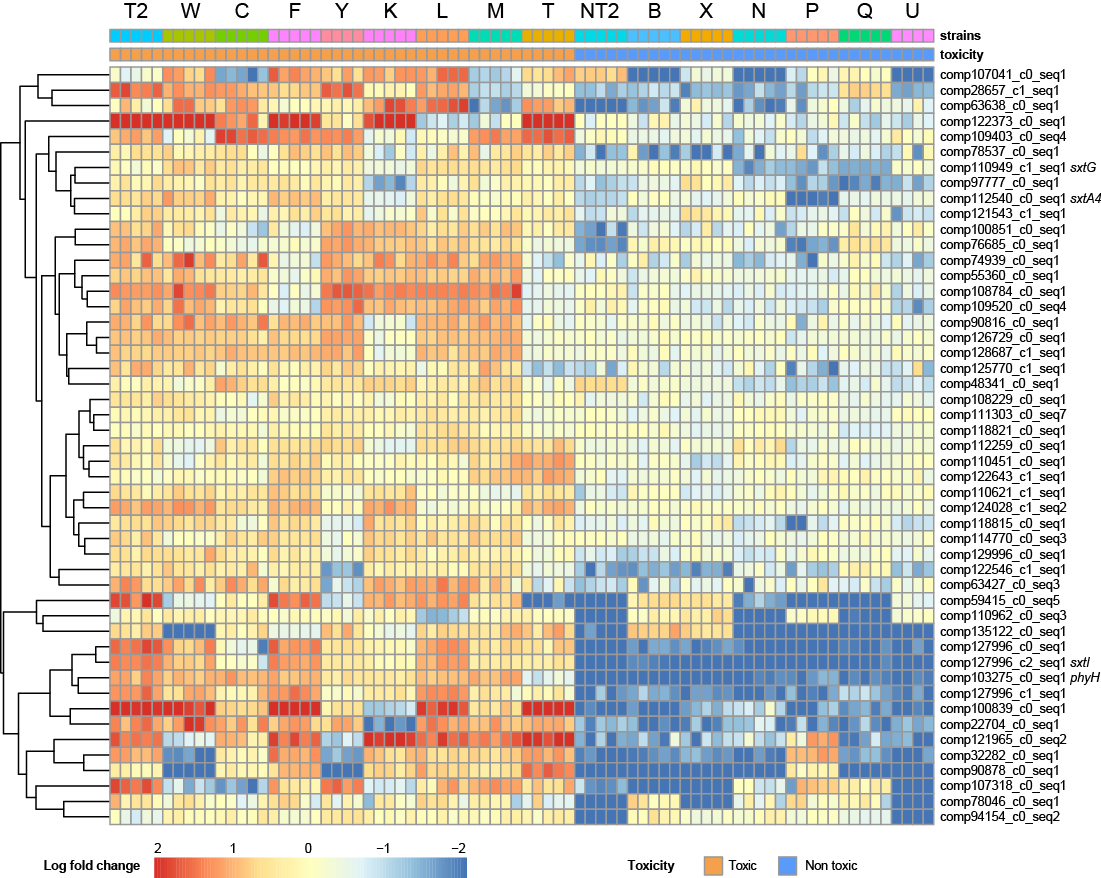

**Supplementary fig. 3:** Expression matrix computed with the linear model, showing overexpressed transcripts in **non-toxic** strains (all of the transcripts could not be represented, thus the results were filtered with a p-value threshold of 0.001). The blue to red scale indicates log2 fold change values between -2 (blue) and 2 (red) (i.e. Blue = underexpressed, red = overexpressed). Log fold change has been set between -2 and 2 for visualization purposes.

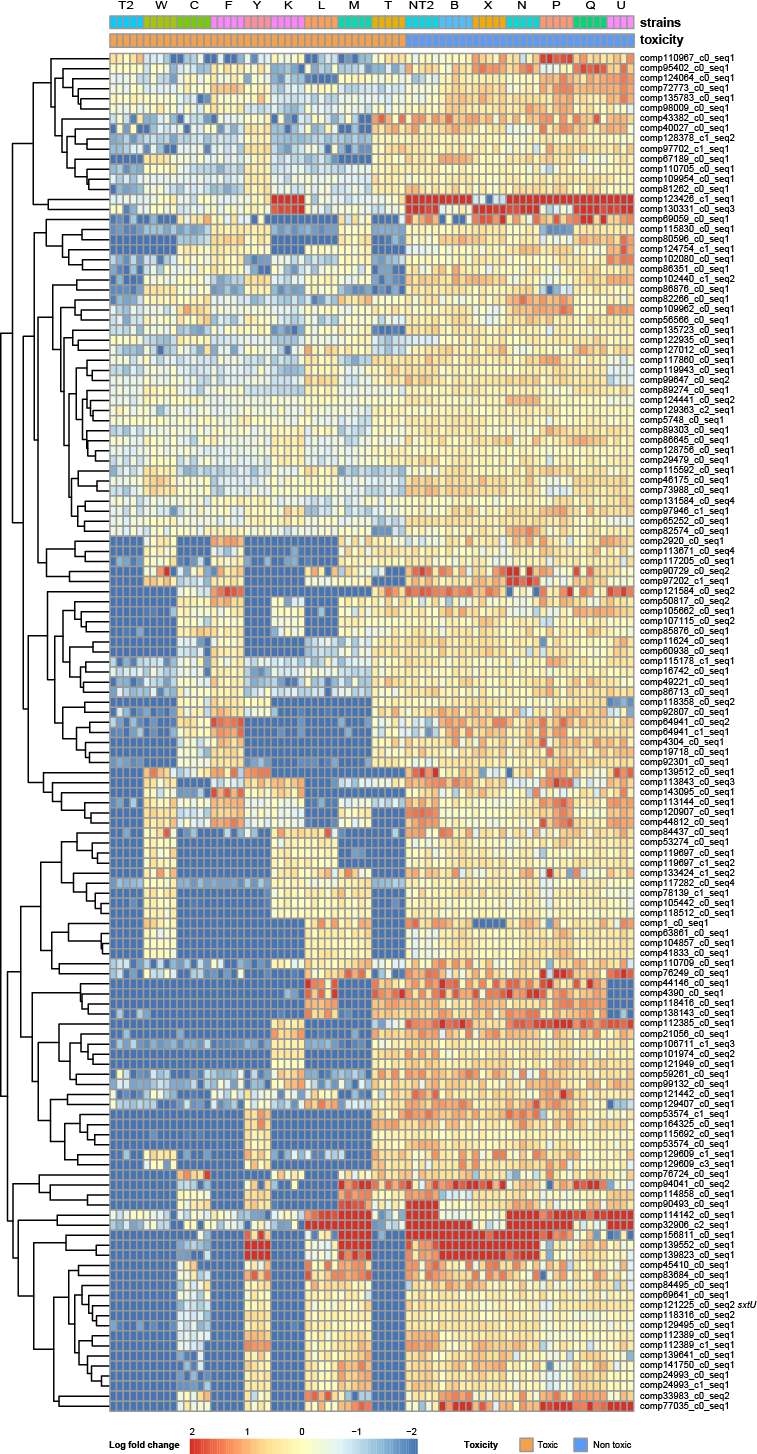

**Supplementary fig. 4:** Expression matrix computed with the linear model, showing overexpressed metabolites in **toxic** strains (all of the features could not be represented, thus the results were filtered with a p-value threshold of 0.01). The blue to red scale indicates log2 fold change values between -2 (blue) and 2 (red) (i.e. Blue = underexpressed, red = overexpressed). Log fold change has been set between -2 and 2 for visualization purposes.

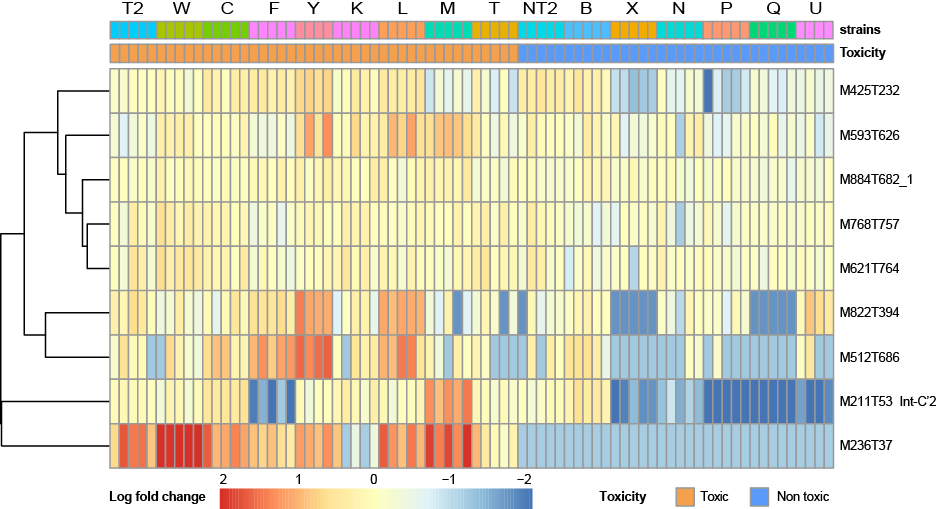

**Supplementary fig. 5:** Expression matrix computed with the linear model, showing overexpressed metabolites in **non-toxic** strains (all of the features could not be represented, thus the results were filtered with a p-value threshold of 0.01). The blue to red scale indicates log2 fold change values between -2 (blue) and 2 (red) (i.e. Blue = underexpressed, red = overexpressed). Log fold change has been set between -2 and 2 for visualization purposes.

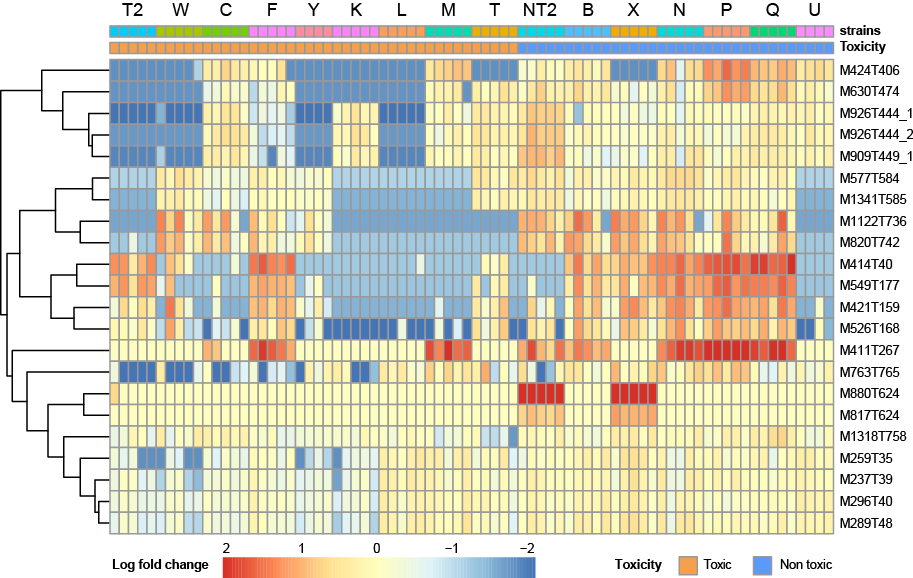

**Supplementary fig. 6:** Boxplots showing the median log-normalized values for features annotated as putative PST biosynthetic intermediates in toxic and non-toxic strains. **(A)** M209T36_2 (putative Cyclic-C’ or dd-doSTX, annotation level 4), **(B)** M225T32 (12ꞵ-d-doSTX, annotation level 2), and **(C)** M241T32 (putative doSTX or 12-β-d-dcSTX, annotation level 4). Normalized values were not significantly (n.s.) different between the two groups for all three features (p-values > 0.05 using non-parametric tests).

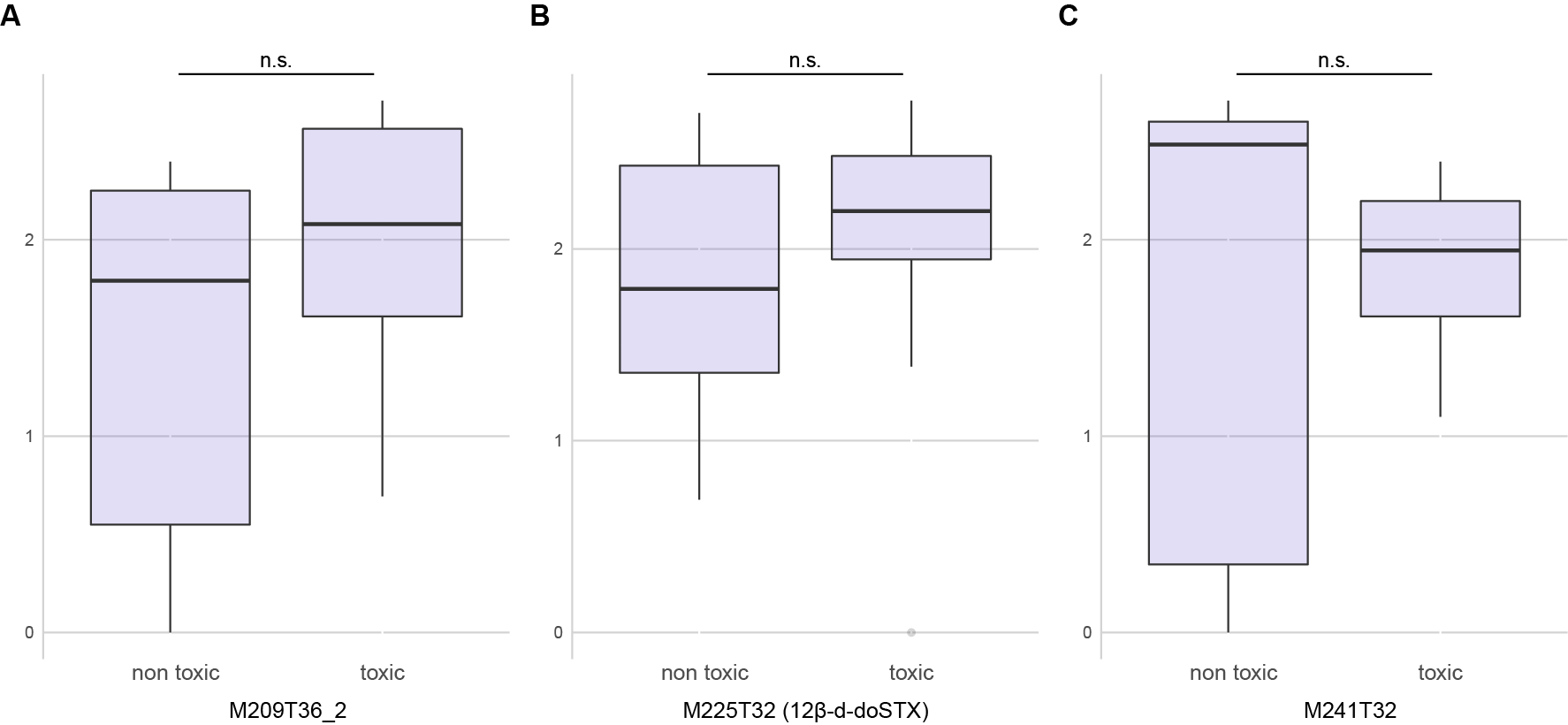

**Supplementary fig. 7**: Blast results (first 12 sequences), represented with bit-scores and % identity, for *A. minutum* **(A)** *sxtI*, **(B)** *sxtG*, **(C)** *sxtA4*, **(D)** PhyH, **(E)** comp118815_c0_seq1 (annotated as Protein Kinase A (PKA)), and **(F)** comp48341_c0_seq1 (annotated as TauD). Sequences of PST-producing dinoflagellates *P. bahamense*, *G. catenatum* and other *Alexandrium* species are shown in orange.

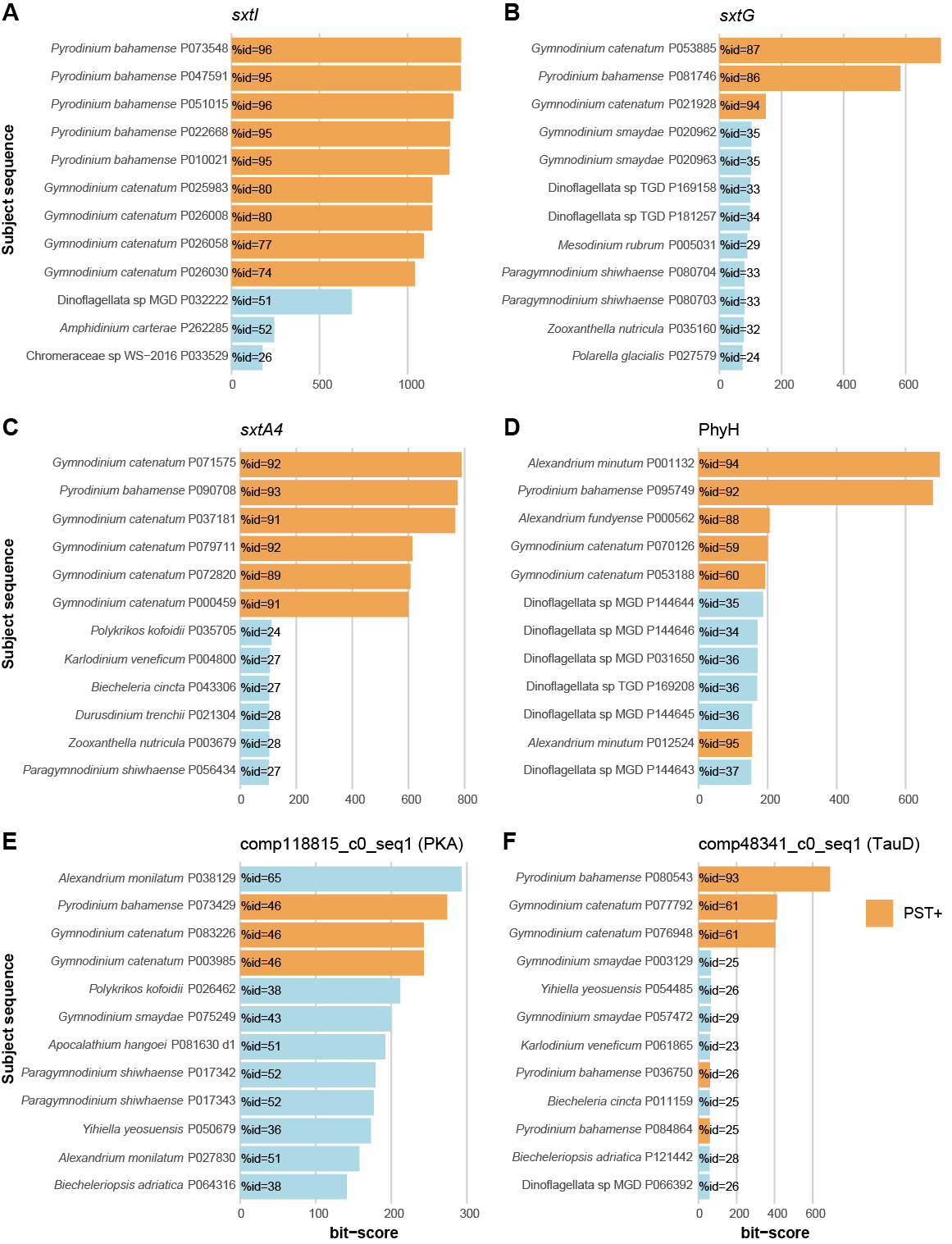

**Supplementary fig. 8:** *A. minutum* putative MFS transporter blast hits (first 12 sequences), represented with bit-scores and % identity, showing sequences from PST-producing (orange) *P. bahamense* and *A. pacificum* as best hits.

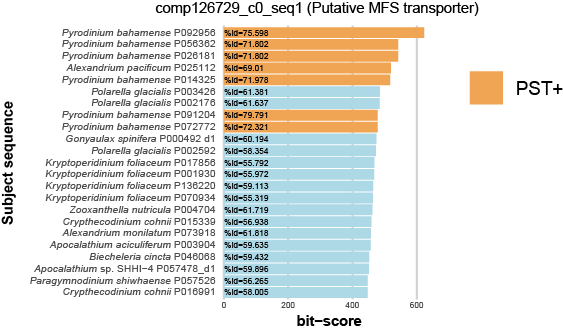
